# Glaucoma and Alzheimer: Neurodegenerative disorders show an adrenergic dysbalance

**DOI:** 10.1101/2022.01.22.477197

**Authors:** Bettina Hohberger, Harald Prüss, Christian Mardin, Robert Lämmer, Johannes Müller, Gerd Wallukat

## Abstract

Glaucoma disease is characterized by an increased intraocular pressure (IOP), glaucomatous alterations of the optic disc and corresponding visual field defects. Even lowering the main risk factor IOP until an individual target level does not prevent this neurodegenerative disorder from proceeding. Several autoimmune mechanisms were discovered, partly showing a functionality. One of these autoimmune phenomena targets the ß2 adrenergic receptor (ß2-AR; i.e. agonistic autoantibodies; ß2-agAAb) and is linked to the elevated IOP and an impaired retinal microcirculation. As neurodegenerative disorder, Alzheimer’s Disease (AD) is postulated to share a common molecular mechanism with glaucoma. In the present study we investigated autoimmune phenomena targeting the ß2-AR in patients with AD. Sera of the patients were analyzed in a rat cardiomyocyte bioassay for the presence of functional autoantibodies against ß2-AR. In addition, different species of amyloid beta (Aß) monomers were tested (Aß1-14, Aß10-25, Aβ10-37 Aß1-40, Aß1-42, Aβ28-40, and [Pyr]-Aß3-42). Our results demonstrate that none of the short-chain Aß (Aß1-14, Aß10-25, or Aβ28-40) showed any agonistic or inhibitory effect on ß2-AR. Contrary, long-chain [Pyr]-Aß3-42, representing a major neurogenic plaque component, exerted an activation that was blocked by the ß2-AR antagonist ICI118.551 indicating that the effect was realized via the ß2-AR. Moreover, the long chain Aß1-40, Aβ1-42, and Aβ10-37 yet not the short-chain Aß peptides prevented the clenbuterol induced desensitization of the ß2-AR. In addition, we identified functional autoantibodies in the sera of AD patients, activating the ß2-AR like the ß2-agAAb found in patients with glaucoma. As autoimmune mechanisms were reportedly involved in the pathogenesis of glaucoma and Alzheimer’s Disease, we postulate that overstimulation of the ß2-AR pathway can induce an adrenergic overdrive, that may play an important role in the multifactorial interplay of neurodegenerative disorders.

## Introduction

Glaucoma disease is a worldwide burden, being the second leading cause of blindness in the industrial nations. This neurodegenerative disease is characterized by an increased intraocular pressure (IOP), a glaucomatous optic disc and visual field defects, going along with disease progression. Up to now the exact pathomechanisms are still elusive. An elevated IOP, the main risk factor, is the target of conservative, laser and surgical therapy. Yet, despite lowering IOP, glaucoma disease proceeds, as no causal therapy is available until now. Neurodegeneration in glaucoma is featured by apoptosis of the retinal ganglion cells (RGC). Immune and autoimmune mechanisms can interplay in this molecular interactions as recent studies suggested. Data of an animal model (mice) yielded a significant RGC loss mediated by T-lymphocytes via heat-shock-proteins in the presence of only less elevated IOP. (Chen et al., 2018) Even normotensive eyes were observed to show an apoptosis of RGC when the contralateral eye was hypertensive (mice model). This effect was attributed to lymphocytes as well.(Gramlich et al., 2020) Apoptosis itself, a vascular dysregulation or other extern factors are able to trigger immune and autoimmune mechanisms. Thus, it is not surprising that diverse autoantibodies were detected in sera of patients with glaucoma. In this context, autoantibodies with a functional consequence are of special interest. Recent studies presented autoantibodies targeting the adrenergic ß2-receptor (ß2-AR) in sera of patients with open-angle glaucoma. These ß2-autoantibodies activate their target (ß2-agAAb) with consecutive overstimulation and desensitization of the receptor contrary to physiological agents. *In vivo* data suggested a link of the ß2-agAAb to the increased IOP and impaired retinal microcirculation.(Junemann et al., 2018;Hohberger et al., 2021a) We propose that a chronic overstimulation of the ß2-AR can result in a dysregulated homeostatic adrenergic balance. Receptor desensitization and internalization might be only one of the molecular features (Wallukat, 2002). If this adrenergic disbalance has sufficient ‘power’ on cellular level, clinical characteristics occur in the patients (*imbalanced autonomic theory*; see also (Szentiványi, 1968)). This adrenergic disbalance is certainly not only a local problem (in the eye). We assume that adrenergic alterations might be even present in extraocular tissue of neurodegenerative diseases. Since several years a common pathomechanism of glaucoma and Alzheimer’s Disease (AD) has been suggested. (Mancino et al., 2018;Inyushin et al., 2019;Rawlyk and Chauhan, 2021) Patients with AD show amyloid beta (Aß) depositions being proposed to be the major mediator of neurotoxicity in the brain.(Wyss-Coray, 2006) Aß is assumed to be one crosslink between glaucoma and AD. RGC express Aß (den Haan et al., 2018;Lee et al., 2020). Especially, intravitreal injection of Aß-42 was observed to induce RGC loss (Simons et al., 2021). Aß is generated after cleavage of the amyloid precursor protein (APP). Ni et al. has shown that a ß2-adrenergic stimulation that activate the gamma secretase is involved in the formation of Aß from the APP (Ni et al., 2006). This cleavage can be mediated by vessel endothelial cell enzymes and platelets, especially when platelets contact the endothelium. Any thrombosis can enhance this Aß generation. Functional AAb against the ß2-AR were not only seen in the sera of patients with glaucoma but also in sera of patients with AD symptoms (Wallukat et al., 2018). These ß2-agAAb but also the long change Aß peptides prevent the desensitization of the ß2AR and induce a permanent stimulation of this adrenergic receptor. As ß2-agAAb show a link to an impaired microcirculation (glaucoma) and Aß expression is increased during impaired microcirculation (i.e. during and after thrombosis; AD), we hypothesize that there is a common molecular mechanisms between adrenergic dysfunction (ß2-agAAb; (I)) and Aß (II). The aim of the study was to investigate sera of patients with AD (I) for the presence of autoantibodies against ß2-AR, and (II) the effect of fragments and the complete Aß peptides on the ß2-AR and whose desensitization.

## Material and Methods

### Patients and controls

Sera of 11 subjects with AD (3 female, 8 male, age: 40-87 years) and 18 subjects with primary open-angle glaucoma (7 female, 11 male, age: 42-76 years) were collected for the present study. Glaucoma patients were recruited at the Department of Ophthalmology and Eye Hospital, Friedrich-Alexander-University Erlangen-Nürnberg [Erlangen Glaucoma Registry, ISSN 2191-5008, CS-2011; NTC00494923].(Hohberger et al., 2019) Patients with AD were recruited at the Department of Neurology, Charite’-Universitätsmedizin Berlin. Blood samples were collected, centrifuged and sera were stored at −20°C. All individuals declared informed consent. The study was performed according to the tenets of the Declaration of Helsinki and approved by the ethic committee of the university of Erlangen, the institutional Review Board of Charite’–Universitätsmedizin Berlin and the Max-Delbrück-Centrum for Molecular Medicine, Berlin-Buch (Y 9004/19, 02.11.2020).

### Primary Open-Angle Glaucoma (POAG) Patients

The criteria of POAG were: an open anterior chamber angle, IOP>21 mmHg (confirmed at least once, measured by Goldmann applanation tonometry), and a glaucomatous optic disc, classified after Jonas.(Jonas et al., 1988) Functional loss of perimetric field were measured and had to be confirmed at least once according to the following criteria: (I) at least three adjacent test points having a deviation ≥5 dB and with one test point with a deviation >10 dB lower than normal, or (II) at least two adjacent test points with a deviation ≥10 dB, or (III) at least three adjacent test points with a deviation ≥5 dB abutting the nasal horizontal meridian or (IV) a mean visual field defect of >2.6 dB.

### Cardiomyocyte bioassay

The cardiomyocyte bioassay was done as described previously (Junemann et al., 2018). Cell culture of cardiac myocytes of heart ventricle (1-3 day-old Sprague-Dawley rats) was used. digested with a 0.25% solution of crude porcine trypsin (Serva, Germany). The cells were dispersed by trypsinization and suspended in a SM20-I medium (Biochrom, Germany), glutamine (Serva, Germany), containing streptomycin (HEFA Pharma; Germany), penicillin (Heyl, Germany), hydrocortisone (Merck, Germany), fluorodeoxyuridine (Serva, Germany), and 10% heat-inactivated neonatal calf serum (Gibco, Germany). The cells were seeded (field density of 160.000 cells/cm^2^) the culture medium was replaced after 24 hours. The cardiomyocytes were cultured for 3-4 days (37°C) before the used in the experiments. Firstly, the basal beating rate of the cardiomyocytes was measured and subsequently the immunoglobulin (Ig) fractions were added for 60 min in a dilution of 1:40. Next, the beating rates (BR) of the cardiomyocytes were counted again on 6 marked spontaneously beating cells or cell cluster (for 15 seconds; at a heated stage of an inverted microscope at 37°C). 1 Unit reflects 4 beats/min (i.e. adrenergic activity). Samples below 2 Units of adrenergic activity were considered as normal. Application of a specific β2-blocker, i.e. ICI 118.551 (0.1μM) was done for identification of the receptor type.

### Preparation of the serum Immunoglobulins

Human IgG were prepared from each patients’ sera by direct ammonium sulfate precipitation (final concentration of 40%; overnight at 4°C). The precipitates were centrifuged (4,000 × g; 30 min) and the pellets were dissolved in dialysis buffer (154 mmol/l NaCl, 10 mmol/l Na_2_HPO_4_/NaH_2_PO_4_, pH 7.2). This procedure was repeated twice. Each sample was dialyzed against phosphate-buffered saline (4°C; 3 days). Storage of the samples was done at −20°C.

### Aß peptides

The fragmented Aß peptides were synthesize by Dr. Beyermann (FMP Berlin). The long change Aß peptides Aß 1-40, Aß 1-42 and also Aß (Pyr) 3-42 were from Sigma-Aldrich.

### Statistical analysis

Statistical analysis was done by SPSS (version 21) and Excel. Dats are shown as absolute value, mean and standards deviation.

## Results

### Patients with glaucoma and Alzheimer’s Disease

Patients with POAG showed a mean beat rate of 4.5±0.1 U (range: 3.3-5.2 U) that was realized via the β2-AR. This increase in the beat rate could be blocked by the specific ß2-blocker ICI 118.551 (Figure 1). These fAAB recognize the second extracellular loop of the β2-AR (Junemann et al., 2018). Functional AAB against the β2-adrenoceptor prepared from patients with AD showed a mean beating rate of 4.5±0.4 U (range: 2.3-7.2 U). As observed for patients with POAG, this increase in the beat rate could be blocked by the specific ß2-blocker ICI 118.551 (Figure 1). Thus, a seropositivity for β2-AAb was observed for patients with POAG and AD. However, the β2-AR fAAB prepared from AD patient sera recognize the first extracellular loop of the β2-AR (Wallukat et al., 2018). Moreover, the activation with the β2-AAb induced in contrast to the classical β2-AR agonists a permanent stimulation of the β2-AR mediated signal cascade without desensitization (Figure 2).

**Fig. 1.**
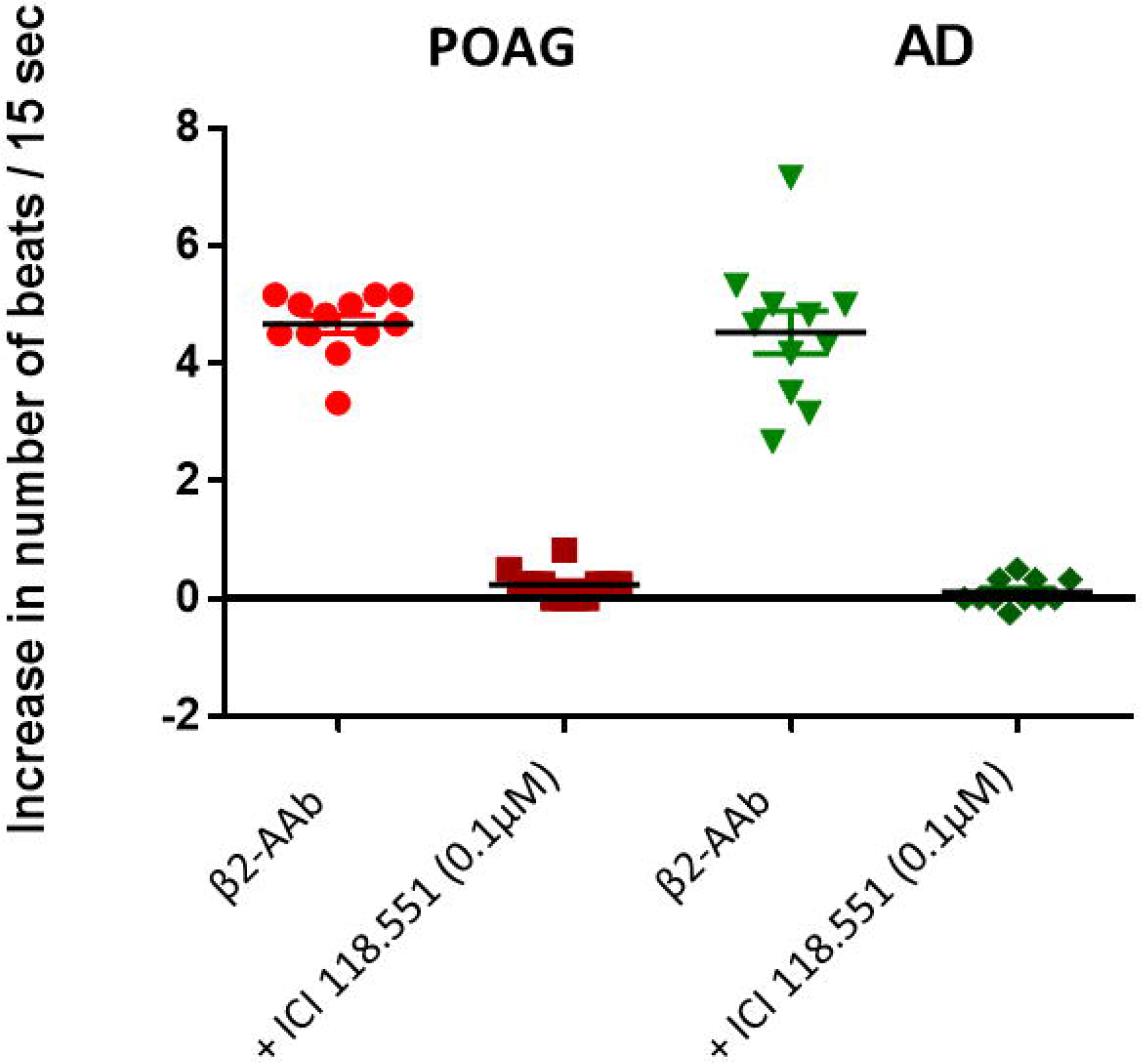
Comparison of the activity of the functional β2-adrenoceptor autoantibodies (β2-AAb) of patients with glaucoma (POAG) and Alzheimer’s disease (AD). Both agonist-like effects were blocked by the β2-adrenoceptor antagonist ICI 118.551 (0.1μM). The experiments with the AAb of the AD patients were done in the presence of the α1-adrenoceptor antagonist urapidil (p<0.05).

**Fig. 2.**
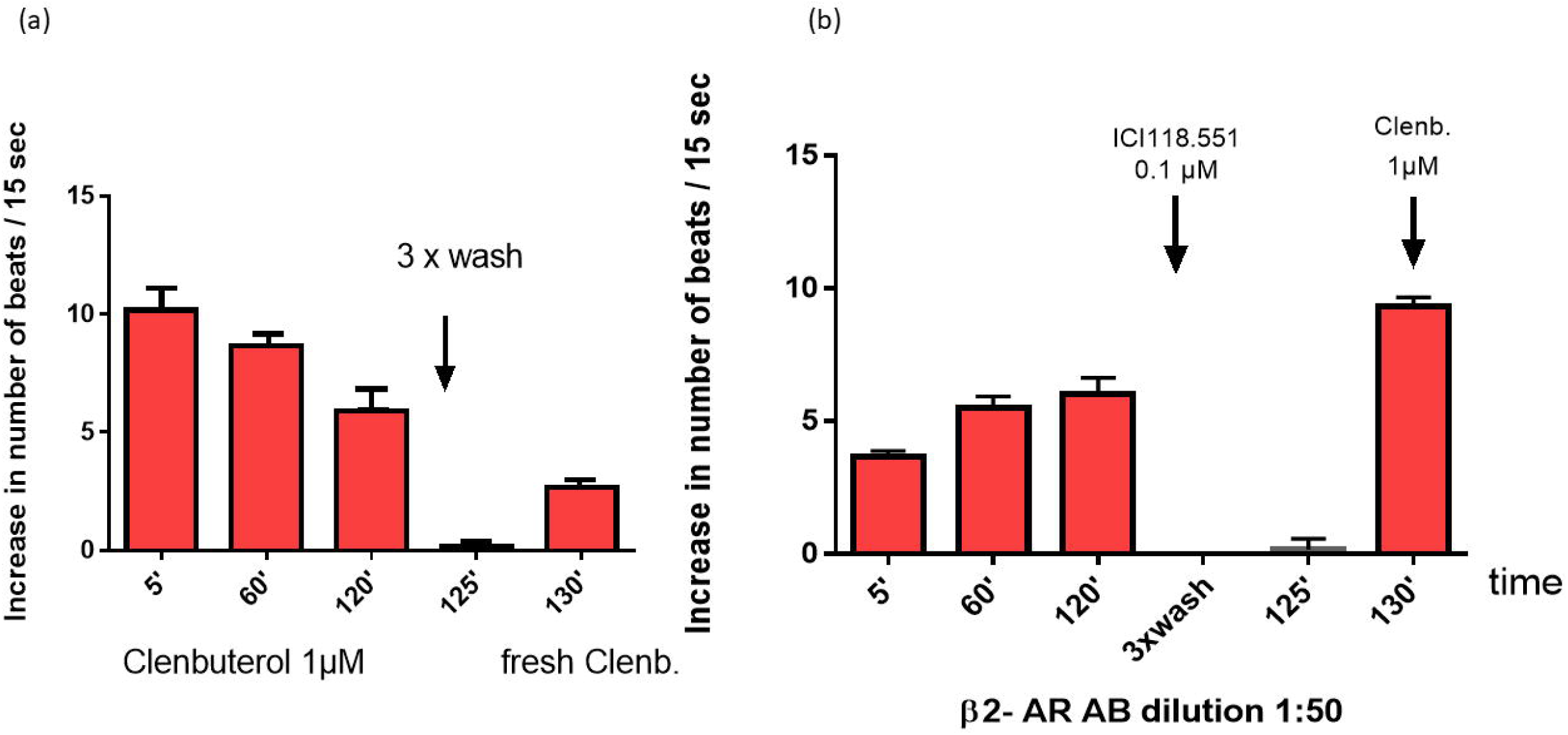
Desensitization of the β2 adrenergic response by the β2-adrenergic agonist clenbuterol (a). This receptor desensitization was missed if the cells were stimulated with the β2-adrenergic AAb prepared from AD patients (b).

### In vitro analysis of amyolid ß (Aβ) peptides effects

In order to identify a possible functional action of Aß peptides on the GPCR different species of amyloid beta (Aß) peptide were tested in the used bioassay. Following Aß peptides were investigated: Aß1-14, Aß10-37, Aß 25-35, Aß 28-40, Aß1-40, Aß1-42, and [Pyr]-Aß3-43). Our results demonstrate that only the Aß peptide [Pyr]-Aß 3-43, but not the other Aß peptides or Aß fragments are able to activate the β2 adrenoceptor similar as the classical β-adrenergic agonists. The long-chain [Pyr]-Aß3-42, representing a major neurogenic plaque component, exerted an activation of the ß2-AR blocked by the ß2-AR antagonist ICI 118.551 but not by the β1-AR antagonists bisoprolol or the α1-AR blocker urapidil (Figure 3). Moreover, the long chain Aß1-40, Aβ 1-42, and Aβ 10-37 but not the short-chain Aß-peptides prevented like the fAAB directed against the β2-AR the clenbuterol induced desensitization of the ß2-AR (Figure 4, table 1).

**Fig. 3.**
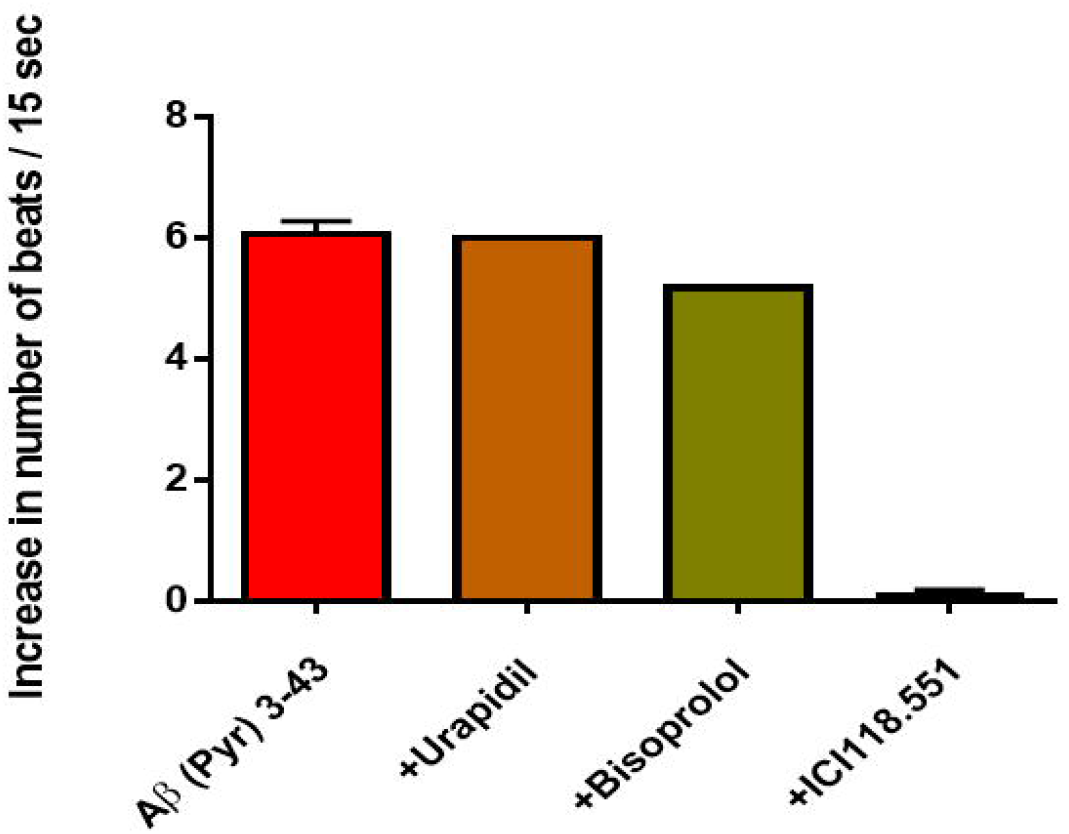
Functional effect realized via the β2 adrenoceptor by the truncated Aβ peptide Aβ [PYR] 3-43: This effect was blocked by ICI 118.551, yet not by the β1 adrenoceptor antagonist bisoprolol or the α1-adrenergic antagonist urapidil.

**Fig. 4.**
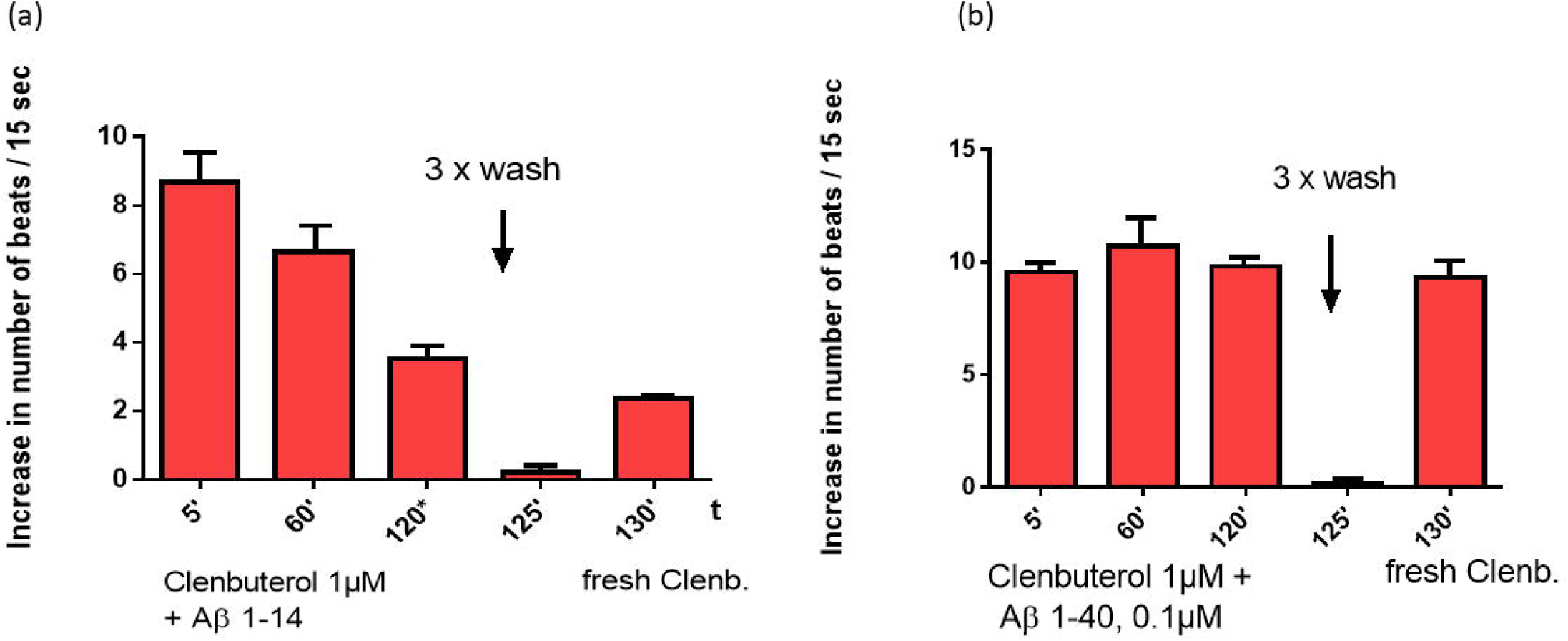
Influence of the Aβ peptides on the clenbuterol induced β2-adrenoceptor desensitization: the desensitization of the β-AR response by clenbuterol, was not influenced by the short chain Aβ peptides (a). However, the long chain Aβpeptides Aβ 10-37, Aβ1-40, and Aβ1-42 prevent the desensitization and exert a permanent stimulation of the β2-AR signal cascade (b).

**Tab.1.** Influence of short and long chain Amyloid β peptides on the clenbuterol induced β2-adrenoceptor desensitization.

| Time |  | 5' | 60' | 120' |  | 125' |  | 130' |
| --- | --- | --- | --- | --- | --- | --- | --- | --- |
| Clenbuterol 1 μM | Clenbuterol 1μM | 10.2±0.3 | 8.7±0.3 | 5.9±0.3 | 3 x wash | 0.2±0.1 | Clenbuterol 1 μM | 2.7±0.1 |
| | + Aβ 1-14 | 8.7±0.6 | 6.7±0.5 | 3.5±0.5 | | 0.2±0.1 | | 2.4±0.1 |
| | + Aβ 25-35 | 8.9±1.2 | 7.4±1.0 | 5.7±0.6 | | 0.2±0.1 | | 2.9±0.2 |
| | + Aβ 28-40 | 10.5±0.2 | 7.8±0.6 | 4.3±0.6 | | 0.3±0.2 | | 2.3±0.2 |
| | + Aβ 1-40 | 10.8±0.2 | 10.7±0.4 | 9.8±0.2 | | 0.6±0.1 | | 9.3±2.9 |
| | + Aβ 1-42 | 8.9±0.1 | 9.3±0.4 | 8.9±0.4 | | 0.2±0.1 | | 9.6±0.3 |
| | + Aβ 10-37 | 9.6±0.2 | 10.1±0.3 | 8.2±0.8 | | 0.4±0.1 | | 10.3±0.2 |

## Discussion

Glaucoma disease is one of the leading causes of irreversible blindness in the developing countries. Its complex pathogenesis is known to be multifactorial with an immune and autoimmune involvement. Clinical studies observed that this neurodegenerative ocular disorder shows several common features with neurodegenerative disorders of the brain (e.g. AD).(Criscuolo et al., 2017;Ramirez et al., 2017;Mancino et al., 2018;Rawlyk and Chauhan, 2021) Thus, it is commonly accepted that both disease entities seem to have common molecular pathways and clinical characteristics. To the best of our knowledge the present study shows a common pathogenic involvement of the β2-adrenergic pathway both in AD and glaucoma for the first time. In sera of patients with AD and glaucoma a seropositivity for ß2-agAAb was observed, respectively. In addition, the long-chain [Pyr]-Aß3-42, representing the major neurogenic plaque component, showed a direct activation of the ß2-AR that was blocked by the specific antagonist ICI118.551. This N-terminal truncated isoform of Aβ can trigger neurodegeneration in a transgenic mouse model (Wirths,2009) and developed a high neurotoxicity (Tekirian,1999). The other functional active Aβ peptides realize their effect in the used test model not via the β2 adrenoceptor. Another aspect of the effects of the Aβ peptides exist in fact that the long chain Aß1-40, Aβ1-42, Aβ28-40 peptides prevented the clenbuterol induced desensitization of the ß2-AR similar as the β2-adrenoceptor autoantibodies. All three mechanisms result finally in a boosted activation and permanent stimulation of the ß2-adrenergic pathways, potentially causing a dangerous adrenergic overdrive realized via the β2-AR.

Glaucoma and AD disease were observed to share common features clinically and biochemically. Patients with open-angle glaucoma are at higher risk (Lin et al., 2014) and showed a higher incidence of AD (Moon et al., 2018). Vis-a-vis patients with AD are at higher risk of glaucoma (Lai et al., 2017). Both, glaucoma and AD, were observed to be age-related disorders (Klein et al., 1995;Molinuevo and Casado-Naranjo, 2014;Doucette et al., 2015;Guerreiro and Bras, 2015;McMonnies, 2017). As the eye, especially retinal tissue, represents “cerebral tissue” considering embryology, it is not remarkable that neurodegeneration and amyloid-β plaques were seen within the retina in AD patients (Whitehouse et al., 1981;Hinton et al., 1986;van de Nes et al., 2008). Degeneration of retinal ganglion cells (RGC) locally and retrograde degeneration of axons in the lateral geniculate body are two findings, being present in AD patients (Hinton et al., 1986;Sadun and Bassi, 1990;Braak and Braak, 1997;Simons et al., 2021). Even amyloid-ß can induce RGC loss within the retina itself (Simons et al., 2021).These neurodegenerative features were also observed in glaucomatous eyes (Lee et al., 2014;Russo et al., 2016;Schmidt et al., 2018;Kosior-Jarecka et al., 2020). The diagnostic correlate of RGC loss can be seen in retinal nerve fiber layer thinning (RNFL) *in vivo*, visualized by magnet resonance tomography and optical coherence tomography (OCT) (Schmidt et al., 2018;Kosior-Jarecka et al., 2020). Both, patients with glaucoma and AD, show a RNFL thinning especially in the superior and/or inferior retina (Blanks et al., 1996a;Blanks et al., 1996b;Sehi et al., 2012;Schrems et al., 2017). Thus, it can be hypothesized that there is common pathway, linking both neurodegenerative disorders (glaucoma and AD).

The autonomic system is a main actor in human body. It regulates important function and is the basis of the human live. Any alterations in the sympathetic and/or parasympathetic pathways results in an impairment or overaction of the autonomic system. Adrenergic receptors are also present on pericytes, thus it can be hypothesized that any alterations in adrenergic activity might influence microcirculation. It is commonly accepted that an impaired microcirculation can induced amyloid plaques in AD (Ishimaru et al., 1996;Armstrong, 2006;Zhang et al., 2020) and is a very early pathogenetic alteration in glaucoma disease (Grieshaber et al., 2007;Wey et al., 2019;Hohberger et al., 2021b). Up to now, it is still elusive which factors induce this impaired microcirculation in AD. Interestingly, recent data showed that an impaired microcirculation is able to change the amyloid-β plaques in a mouse model (Zhang et al., 2020). Even in glaucoma pathogenesis, the exact pathogenesis is under investigation. There is evidence that an involvement of the autonomic system, especially the adrenergic ß2-mediated pathway might be present in dysregulation of IOP (glaucoma) (Hohberger et al., 2020;Hohberger et al., 2021c) and capillary microcirculation (glaucoma (Hohberger et al., 2021a) and AD), as patients with glaucoma and AD showed a seropositivity for ß2-AAb, respectively.

It was shown that patients with AD develop functional active autoantibodies directed against the ß2-AR. These ß2-agAAb activate the ß2-AR in a similar manner as the ß2 adrenergic agonists. (Wallukat et al., 2018) but in contrast to the agonists the ß2-agAAb prevent the desensitization of the corresponding receptor. This observation was also made in glaucoma patients (Junemann et al., 2018). The agonist like effect of this ß2-agaAAb was blocked by ICI118.551, yet not by the ß-AR antagonist bisoprolol. Moreover, the agonistic effect of the ß2-agAAb was neutralized by a peptide corresponding to the second extracellular loop of this receptor. In contrast to the classical ß-AR agonists, ß2-agAAb activate the ß2-AR mediated signal cascade permanently. Moreover, the ß2-agAAb also prevent the desensitization, normally seen for the classical ß2-AR agonists, when agonists were added to ß2-agAAb prestimulated cardiomyocytes. This permanent stimulation without desensitization of the β2-AR may represent a pathogenic factor that played an essential role in the pathogenesis of several diseases (Wallukat et al., 1991). Another feature of these ß2-agAAb is the possible antagonistic action of these ß2-agAAb directed against the β2-AR. It was shown by Mijares et al. that monovalent Fab fragments of monoclonal β2-agAbbs exhibit antagonistic and not agonistic properties (Mijares et al., 2000). However, if these monovalent Fab fragments were crosslinked by an anti Fab antibody than the agonist-like effects were restored (Wallukat and Schimke, 2014). These data demonstrate that the ß2-agAAb also can act in an antagonistic manner under particular circumstandes.

In patients with AD disease the ß2-adrenergic dysregulation might be mediated in three different ways. First is the direct activation of the ß2-AR by its classical natural agonist adrenaline or synthetic compound like isoprenaline. Both compounds activate the ß1-, the ß2- and the ß3-AR. The stimulation with adrenaline, isoprenaline or the more ß2-AR specific agonist clenbuterol induced a positive chronotropic response in the cardiomyocyte bioassay, which could be blocked by the specific ß2-AR antagonist ICI118.551. Furthermore, clenbuterol caused like the most agonists during a permanent treatment a desensitization of the of the ß2-AR mediated response. After an incubation time of two hours, a washing procedure and a new stimulation with the same agonist concentration the cardiomyocytes respond only with a third of maximal response (Fig. 2a).

Secondly, it was shown that the activation of the ß2-AR may play a role in the formation of amyloid β (Aβ). ß2-AR agonists activate the gamma-secretase, forming Aβ from the amyloid precursor proteins (APP), resulting in an elevated accumulation of Aβ (Ni et al., 2006). Moreover, a micro perfusion of Aβ1-40 in combination with the ß2-adrenergic agonist clenbuterol into the hippocampus caused an augmented degradation of the Aβ1:40 peptide in comparison to the controls without clenbuterol (Richter R.M, 2010). Some of these degradation products of the Aβ1-40 and Aβ1-40 itself can stimulate different G-Protein coupled receptors (GPCR) in a similar manner as the classical receptor agonists. For example, the Aβ-fragments Aβ25-35 and Aβ10-37 activate the α1-AR and this activation was abolished by the antagonist prazosin or urapidil. These Aβ-peptides recognize the first extracellular loop of the α1-adrenergic receptor and the Aβ25-35 and Aβ10-37 induced activation could be blocked by peptides corresponding to the first extracellular loop (Haase et al., 2013). Such α1-adrenoceptor antibodies can influence moderately the contractility of the coronary and middle cerebral artery and strongly the renal artery of the rat (Yan et al., 2009). Moreover, Tohda et al. demonstrated that Aβ 25-35 induced memory impairment, axonal atrophy and synaptic loss (Tohda et al., 2004).

Wang and coworker (a,b) observed in prefrontal cortical neurons that Aβ1-42 can bind to the β2-AR, and activate the protein kinase A (PKA) in these neurons (Wang et al., 2010;Wang et al., 2011). This observation was not confirmed in our experiments using a functional cardiomyocyte assay. In our bioassay Aβ 1-40, Aβ1-42 but also Aβ1-14 activate the fast estradiol receptor GPR 30 (unpublished data). In contrast, the N-terminal truncated Aβ peptide [Pyr] 3-43 that representing a major neuritic plaque component was able to activate the β2-AR subtype in our cardiomyocyte model. This Aβ peptide, activate the ß2-AR in similar manner as the agonist clenbuterol. This activation was neither influenced by the α1-adrenergic antagonist urapidil nor the β1-antagonist bisoprolol. Only ICI118.551 was able to block the activity induced by Aβ [Pyr] 3-43. Therefore, we assume that in the experiments of Wang on neuronal tissues the Aβ-peptides Aβ1-40 and Aβ1-42 were truncated to Aβ [Pyr] 3-43 by endogenous peptidases that can recognize the β2-AR subtype.

A third possibility to influence the β2-AR activity can be seen in fact that the long chain Aβ peptides Aβ1-42, Aβ1-40, and Aβ10-37, yet not the short breaking products of the amyloid Aβ1-14, Aβ25-35 and Aβ28-40 prevent desensitization of the β2-adrenergic agonist clenbuterol. The combination of the long chain Aβ peptides and clenbuterol induced a permanent activation of the β2-signal cascade similar to the response seen for the agonist-like β2-AAbs. This observation indicates that the long chain Aβ can also contribute on the adrenergic overdrive.

We hypothesize that a dysregulated β2-mediated signal cascade might be a molecular mechanism of an autonomic dysregulation, both observed in neurodegenerative disorders, e.g. glaucoma and AD. A seropositivity for β2-AAb and the β-adrenergic agonists itself can enhance β2-AR activation. Additionally, to β2-AAb and the Aβ peptide Aβ [Pyr] 3-43 were also able to activate this signal cascade directly. Furthermore, the long chain Aβ peptides induce like the β2-AAb a permanent stimulation and deactivate the protection mechanism of the cells against an adrenergic overstimulation. All these mechanisms may play an important role in the pathogenesis of Glaucoma and AD.

## Conclusion

Glaucoma and AD disorder are known to be neurodegenerative disorders with an involvement of autoimmune mechanisms and impaired microcirculation. As both disease entities showed a seropositivity for β2-AAb we postulate that an overstimulated ß2-AR pathway can induce an adrenergic overdrive, which can be enhanced by amyloid-β peptides in AD patients.

